# Unaffected functional recovery after spinal cord contusions at different circadian times

**DOI:** 10.1101/2021.03.30.437713

**Authors:** Lukasz P. Slomnicki, George Wei, Darlene A. Burke, Scott R. Whittemore, Sujata Saraswat Ohri, Michal Hetman

**Author notes:** Corresponding author: Michal Hetman, M.D., Ph.D., Kentucky Spinal Cord Injury Research Center, Departments of Neurological Surgery, Anatomical Sciences & Neurobiology, and Pharmacology & Toxicology, 511 S. Floyd St., MDR616, University of Louisville School of Medicine, Louisville, KY, 40292, / /.

## Abstract

The circadian rhythms of gene expression drive diurnal oscillations of physiological processes that determine the acute injury response including immunity, inflammation and hemostasis. While outcomes of various acute injuries are affected by the time of day at which the original insult occurred, such diurnal influences on recovery after spinal cord injury (SCI) are unknown. We report that several key regulators of circadian gene expression are differentially expressed in uninjured spinal cord tissue of naïve mice at Zeitgeber time 1 (ZT1) or ZT12, where ZT0 or ZT12 are times when lights are turned on or off, respectively. However, mice that received moderate, T9 contusive SCI at ZT0 or ZT12 showed similar recovery of locomotion as determined using the ladder walking test and the Basso mouse scale (BMS) over a 6 week post-injury period. Consistent with those findings, terminal histological analysis revealed no significant differences in white matter sparing at the injury epicenter. Therefore, locomotor recovery after thoracic contusive SCI is not affected by the time of day at which the neurotrauma occurred at least when comparing the beginning to the end of the mouse active period.

## Introduction

Time of day affects the incidence of various acute pathologies including myocardial infarct (MI) and ischemic stroke [1]. Moreover, the severity of those injuries may also be influenced by the time of day at which they occur [2, 3]. Such effects stem from circadian rhythmicity of biological processes that determine risk of a blood vessel occlusion and/or rupture and/or modify the tissue injury response [1]. Thus, in morning hours when the human active period begins, circadian maxima (acrophases) of blood pressure, sympathetic tone, and hemostasis may explain higher occurrence and greater severity of MI and stroke at that time [1]. In addition, circadian modulation of metabolism, pro-inflammatory potential, immunity, anti-oxidant defenses and blood-tissue barriers may directly affect sensitivity to acute injuries [1, 4–6].

Circadian rhythmicity of biological processes is mediated by oscillations of gene expression produced by a set of conserved transcription factors (TFs) of the clock pathway including BMAL1, CLOCK and NPAS2 [7]. The clock pathway in hypothalamic suprachiasmatic nucleus (SCN) neurons synchronizes clock pathways in other cells of the body. The clock pathway TFs engage feedback loops that underlie oscillating expression of them and their regulators.

In rodents, the activity of the effector outputs of the clock pathway in most non-SCN tissues is low at ZT18-0 (late night/early morning) or high at ZT6-12 (afternoon/early evening), respectively [8] [9, 10]. Such oscillations coincide with differential responses to such challenges as MI, infection, endotoxic shock or autoantigen exposure while genetic disruption of clock signaling nullifies those time of day effects [11–15].

Time of day effects are also documented in several models of acute brain injury [16–21]. However, the maximum severity of brain damage peaked at distinctly different times dependent on the model used [16–21]. Such variability suggests that unique, injury-specific pathogenic mechanisms may be differentially sensitive to circadian regulation. There are no reports of circadian effects on spinal cord injury (SCI).

BMAL1 is the principal non-redundant TF of the clock pathway output [7]. After moderate contusive SCI at the T9 level, *Bmal1*^−/−^ mice showed enhanced locomotor recovery, increased white matter sparing as well as reduced inflammation and improved blood-spinal cord barrier function in the injury epicenter region [22]. Therefore, the pathogenesis of SCI may be regulated by circadian rhythms. The current study was initiated to test whether SCI outcomes differ if the injury occurs at the beginning or end of the day, when the mouse active period ends or begins, respectively.

## Methods

### Animals

Six-week old C57Bl/6 wild-type female mice were obtained from the Jackson Laboratory (Bar Harbor, ME). Animals were maintained in a 12:12 light-dark cycle (6:00 light on, 18:00 light off) with food and water available *ad libitum* for 2 weeks. After five days of habituation to handling (performed in the same room where behavioral assessments were later performed) mice were randomly assigned to different experimental groups. All animal procedures were approved by the University of Louisville Institutional Animal Care and Use Committee and strictly adhered to NIH guidelines.

### Spinal cord injury

Avertin anesthesia, T9 spinal cord contusion (50 kdyn, IH impactor /Infinite Horizons, Lexington, KY/) and post-surgery care were performed as previously described [22, 23] (see Supplementary Methods for detailed information including anesthesia and post-surgery analgesia). The surgeries were performed at ZT0-1.5 (6:00-7:30, n=11) or ZT12-13.5 (18:00-19:30, n=11) by the same team of investigators 12 h apart. Both groups were given identical post-surgical care and maintained under the same conditions for six weeks. Three mice were lost (1 in ZT0 euthanized after accidental rupture of the bladder during bladder expression, 2 in ZT12 were found dead at dpi 8 and dpi 10). See Supplementary Table 1 for detailed contusion parameters (recorded force, displacement, velocity).

### Assessments of locomotor function

All behavioral assessments were performed at the same time for both groups of mice by individuals without knowledge of group assignment. Hindlimb locomotor function was evaluated in an open field using the Basso Mouse Scale (BMS) by raters trained by Dr. Basso and colleagues at the Ohio State University [24]. Evaluations were performed weekly, first before the injury to determine baseline values, and then for six weeks starting at week 1 after SCI. The horizontal ladder test was performed as described previously using Columbus Instruments Sensor and RS232 Mini Counter (Columbus Instruments; Columbus, OH, USA, 2.5 mm rungs spaced 3.5 cm apart) [25]. Briefly, each animal underwent five stepping trials per session and the total number of footfalls was quantified for the left and right limbs, respectively. A baseline session before SCI was followed by bi-weekly assessments starting at 2 weeks post-injury.

### White matter sparing

was performed as described previously [23]. Briefly, after completion of behavioral analyses (day post-injury 42), mice were deeply anaesthetized and perfused transcardially with ice cold phosphate buffered saline (PBS) and then 4% paraformaldehyde (PFA) in PBS. Twenty μm serial transverse cryosections from a 4 mm spinal cord segment centered at the injury epicenter were stained for myelin with iron-eriochrome cyanine (EC). For each animal, the section with least amount of myelinated white matter was identified as the injury epicenter. White matter sparing was defined as % relative white matter area (per total section area) at the epicenter as compared to the relative white matter area 2 mm rostral from the injury epicenter.

### qRT-PCR analysis of circadian oscillations of gene expression

Naïve mice (coming from the same batch of animals as that used for SCI and undergoing same handling habituation) were deeply anesthetized and transcardially perfused with ice cold PBS at ZT1 or ZT12 to collect a 5 mm segment of the thoracic spinal cord and the liver. Total RNA extraction, synthesis of cDNA and SYBR Green-based qPCR analysis using the ΔΔCt quantification method and *Gapdh* as a normalizer followed previously described methodology [22]. See Supplementary Table 2 for primer information. The results were compared to publicly available data on oscillations of the clock pathway mRNAs in various tissues of 7-8 week old C57Bl6 mice including the brain stem, the cerebellum, and the liver (http://circadb.hogeneschlab.org/mouse) [8].

### Statistical analyses

Repeated measures ANOVA (RM ANOVA) followed by Tukey *post hoc* tests was used for analyzing BMS and horizontal ladder locomotor recovery data.

Gene expression and white matter sparing data were analyzed using the non-parametric Mann-Whitney *u*-test.

## Results

In various tissues including intact male or female rat spinal cord as well as male C57Bl6 mouse brain stem, cerebellum or liver transcript levels for most clock pathway mRNAs oscillate with a maximum amplitude at or around ZT0 and ZT12 [26] [8] (http://circadb.hogeneschlab.org/mouse, Supplementary Fig. S1 and S2). Such oscillations indicate activity of the clock pathway, as its core components are also clock pathway-regulated at the transcriptional level [7]. Therefore, levels of selected clock pathway transcripts were analyzed at ZT1 and ZT12 in the intact spinal cord and the liver of naïve mice taken from the same cohort that was used for SCI studies. At ZT12, *Bmal1* decreased by 42.5% in comparison to ZT1 (Fig. 1A). Consistent with increasing BMAL1 TF activity during the rodent inactive period, expression of several BMAL1 target genes including *Nr1d1, Nr1d2, Cry1, Per1, Per2, Per3,* and *Dbp* was higher by 25-65% at ZT12. At that time, *Bmal1* showed a 98% decline in the liver with several of its target genes showing strong increases by 50-90% (Fig. 1B). These data validate natural modulation of the clock pathway in the spinal cord and liver tissues of mice that were used for SCI experiments. The findings are consistent with greater circadian regulation of the transcriptome in the liver as compared to the CNS [8] and the reported maximum amplitude for many clock pathway genes at the start and the end of the mouse active period (Supplementary Fig. S1 and S2).

**Figure 1.**
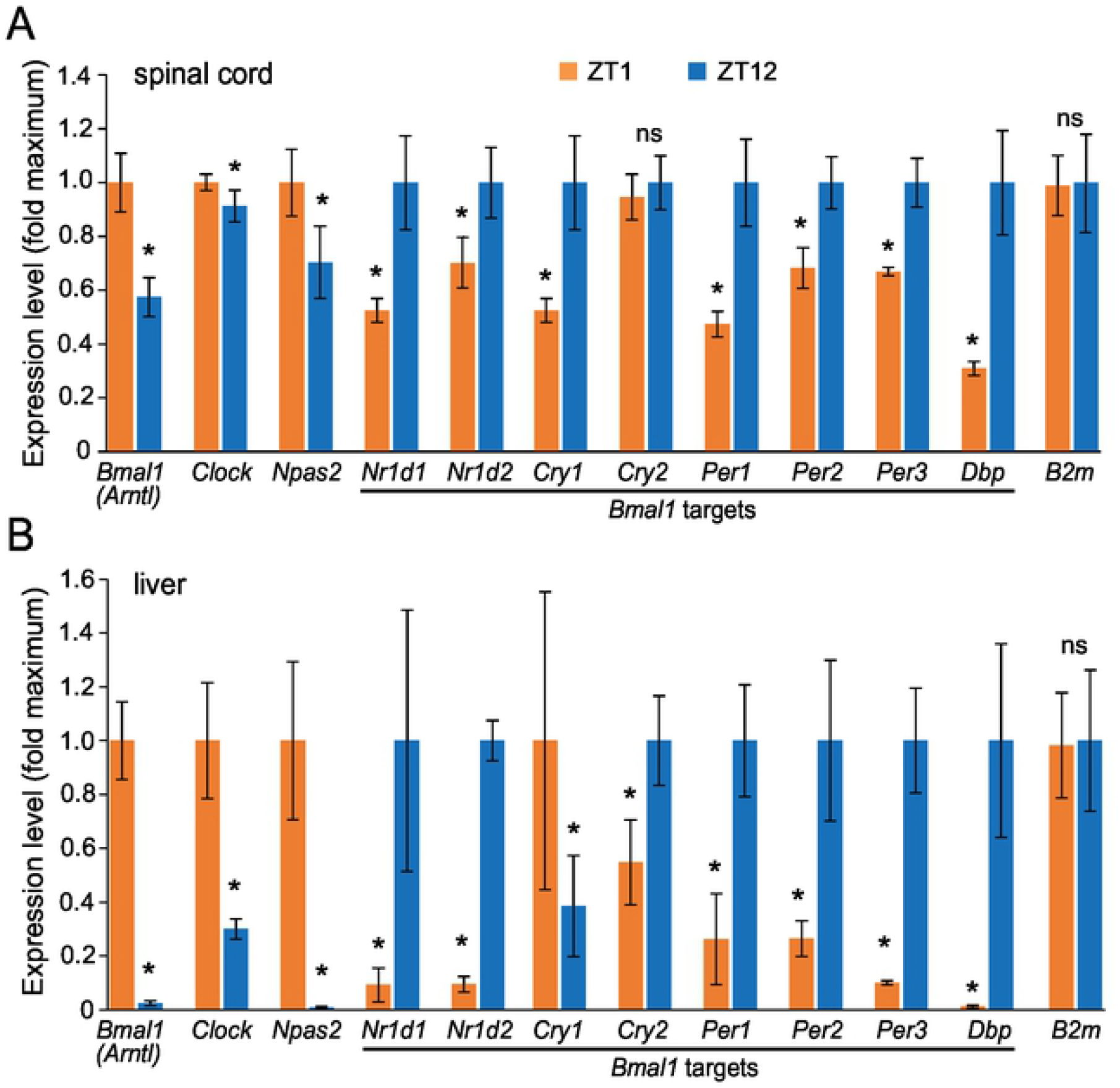
Circadian effects on expression of clock pathway mRNAs in the intact mouse spinal cord. Levels of mRNAs were determined at ZT1 and ZT12 by qPCR using total RNA from the lower thoracic segment of the spinal cord (A) and the liver (B). *Gapdh* was used as a normalizer for expression level determinations, *B2m* was also included as an additional normalizing transcript. Note that lower levels of *Bmal1* expression at ZT12 coincide with increased levels of several BMAL1 target genes suggesting increased activity of the clock pathway output. The observed differences in the spinal cord are consistent with reported maximal amplitudes of clock pathways mRNA in other non-SCN regions of mouse brain at the beginning and the end of the active period (Supplementary Fig. S1 and S2). For each transcript, data represent average fold change of a time point with maximal expression ± SD; *, p<0.05, ns>0.05, *u*-test; n=3 mice/time point.

To determine whether oscillations of the clock pathway activity at the time of injury correlate with a long-term locomotor recovery, moderate T9 SCI was performed at ZT0 or ZT12. Mean displacement was similar for both groups (567.3 ± 88.7 at ZT0 or 575.0 ± 83.1 μm at ZT12, p>0.05, *t*-test) suggesting no difference in severity of the primary injury. Similar recovery of hindlimb function was revealed with terminal BMS scores of 4.90 ± 0.61 or 4.83 ± 0.32 and ladder errors of 9.64 ± 3.04 or 8.87 ± 5.41 for ZT0 or ZT12, respectively (Fig. 2). In rodents with low thoracic level contusive SCI, white matter loss at the injury epicenter is the primary determinant of functional deficits [24, 27]. Therefore, the observed lack of altered functional recovery correlated well with the lack of significant differences in % spared white matter between the groups (Fig. 2C, D).

**Figure 2.**
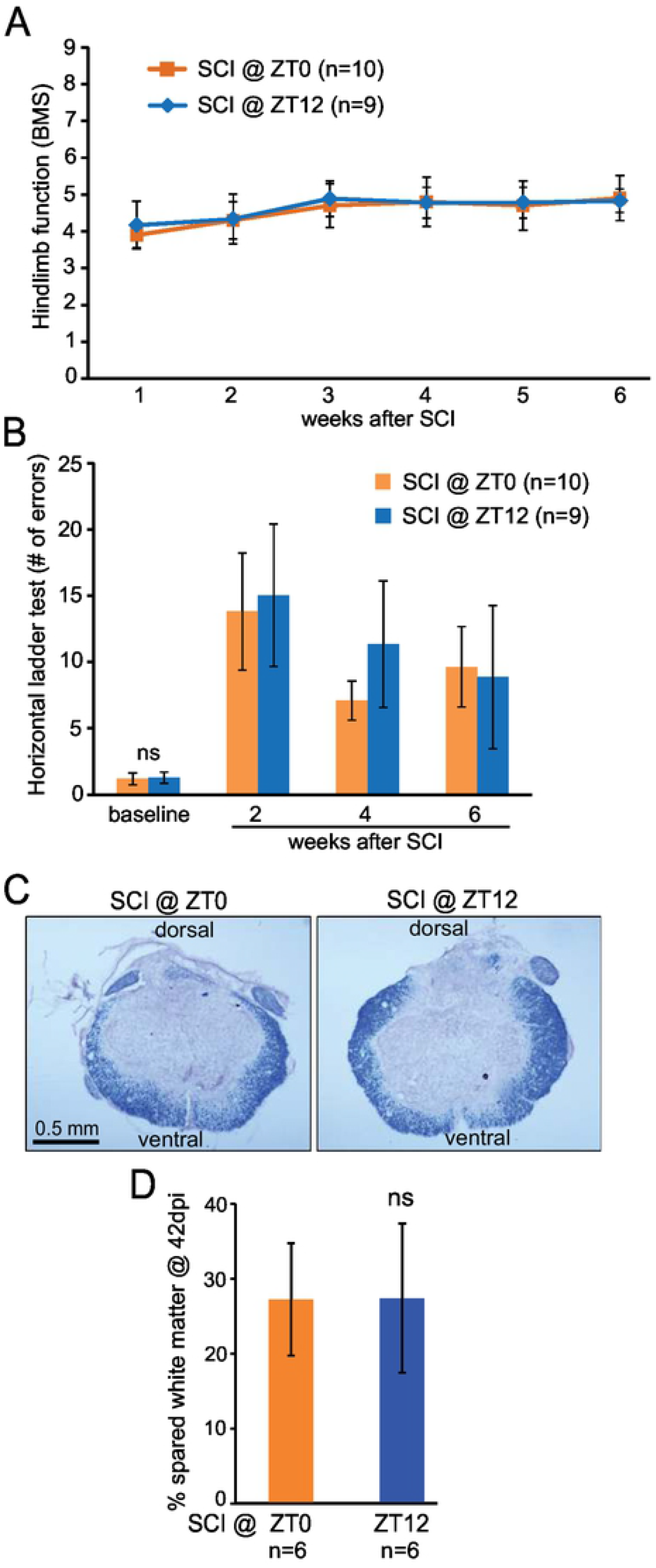
Similar locomotor recovery and white matter sparing after SCI at ZT0 or ZT12. ***A, B,*** Locomotor recovery after T9 50 kdyn IH contusion was quantified using the BMS (A) or the horizontal ladder test (B). The surgeries were done 12 h apart and all the assessments were done at the same time. For either parameter, RM ANOVA showed significant effects of time after injury but not time of injury (time of injury, BMS: F_1,99_=0.64, p=0.43; time of injury, ladder: F_1,46_=1.9, p=0.17; see Supplementary Table 3 for more results). ***C, D,*** WMS was analyzed after completion of behavioral assessments 6 weeks after SCI. ***C,*** Representative images of EC myelin staining in spinal cord sections cut through the injury epicenter. ***D,*** similar % spared white matter in ZT0 SCI and ZT12 SCI mice (ns, p>0.05, *u*-test). Data represent means ± SD.

## Discussion

Our results suggest that after contusive SCI, impairment of hindlimb function and its subsequent recovery is similar irrespective of whether the injury occurred at the beginning or the end of the mouse active period. In addition, consistent with published clock pathway gene expression studies of rodent CNS tissue, we report relatively higher or lower activity of the clock pathway at those two times of day, respectively [8, 9]. As reliable determination of the nadir and the zenith for a circadian-regulated mRNA requires probing at a minimum of 6 different timepoints/day across at least two days, we do not have sufficient data to determine the spinal cord rhythm of clock pathway transcripts [28]. However, we can rely on published reports and/or publicly available data sets from other areas of the mouse CNS as well as other organs that, apart from SCN, show similar phase of oscillations for all major mediators of the clock pathway with a zenith of the clock pathway activity by the start of the active period and a nadir at its end (Supplementary Fig. S1 and S2). In addition, our spinal cord or liver data show that clock pathway mRNA changes at ZT1 and ZT12 are of similar magnitude as maximal amplitudes reported in the mouse CNS or liver, respectively (Fig 1, Supplementary Fig. S1 and S2). One notable exception is *Nr1d1/Nr1d2* which peaks in the middle of the inactive period (Supplementary Fig. S1). Therefore, our analysis likely underestimates its maximal circadian oscillations in the spinal cord and the liver. However, the presented findings confirm that in the cohort of mice that was used for SCI studies, natural circadian oscillations of gene expression occurred in the spinal cord and the liver and that their phase was likely similar to that reported in rodents. Thus, circadian rhythmicity of gene expression and its effector biological processes could be considered as a potential variable that determines outcome of SCI.

While this report is the first to address the question of time of day effects on pathogenesis of SCI, others have documented existence of circadian modulation in several acute CNS injury models. Thus, in a middle cerebral artery occlusion (MCAO) model of rat stroke, acute infarct volume was three times larger with a stroke at ZT22 than at ZT10 [16]. However, relatively moderate circadian effects were reported in a murine MCAO model [21]. Global brain ischemia at ZT14 resulted in maximal hippocampal apoptosis in a rat, while the greatest hippocampal damage followed ZT6 ischemia in mice [17, 19]. In a mouse subarachnoid hemorrhage model, less apoptosis was observed in the hippocampus and the cortex when the insults occurred at ZT12 as compared to ZT2 [20]. Closed skull traumatic brain injury (TBI) in rats at ZT17 reduced brain damage area and acute mortality with a transient improvement in locomotor function as compared to ZT5 TBI [18]. These results indicate that dependent on the CNS injury model used, injuries that occurred in the second half of the active phase through the start of the inactive phase, when clock pathway activity reaches a nadir, produced maximal or minimal pathology at acute/subacute phases. While the current SCI time of day study did not reveal significant differences in long term recovery or tissue sparing, the aforementioned reports of circadian effects focused on acute/subacute pathological changes with long term functional recovery or lesion healing either not examined or unaffected. Hence, in several types of acute CNS injury in rodents, including mouse moderate SCI, the time of insult may have, at most, only transient effects on secondary damage without long lasting functional impact.

One could argue that the severity of the SCI paradigm used for this work may have been too high to detect time of injury effects. This is unlikely as this moderate IH contusion has been used in several mouse studies in which locomotor recovery and terminal white matter sparing were increased or decreased by various genetic or pharmacological manipulations that targeted the secondary injury cascades [22, 23, 29–32]. Hence, excessive primary damage is unlikely to explain our negative findings.

Why is post-SCI recovery not affected by the time of injury despite our prior findings that *Bmal*^−/−^ mice have improved outcome [22]? While there are many possible explanations to reconcile those observations, one should note that BMAL1 has been implicated in gene expression regulation beyond circadian rhythms [33, 34]. Loss of such a regulatory activity was proposed to contribute to a neurological phenotype of BMAL1-deficient mice and may have also played a role in SCI outcome [33]. Moreover, altered clock pathway/BMAL1 activity after, but not before, and/or at the time of SCI may be a critical contributor to secondary injury and long-term recovery after SCI. Interestingly, acute increases of at least some clock pathway components were observed in the injured mouse spinal cord tissue [22]. Those findings may indicate that after SCI, the clock pathway is reset to a new, post-injury time. Hence, modulation of the secondary damage by the injury-regulated activity of the clock pathway may override any earlier influences from natural circadian oscillations. Lastly, a confounding factor that may alter the significance of natural clock pathway oscillations in experimental SCI may be pre- and post-surgery care including such potential clock-resetting stimuli as anesthesia and analgesia [35, 36].

## Conclusions

Current work using moderate contusive SCI indicates no major injury time of day effects on long term locomotor recovery and white matter integrity in mice at least when comparing the beginning to the end of the mouse active period. Obtaining clinical data to test if time of injury affects SCI outcome in humans will be extremely difficult as confounding injury co-morbidities and variable extents of spontaneous recovery will restrict necessary samples sizes [37, 38]. However, future preclinical analyses to address potential effects on additional outcomes such as immune system dysregulation [39] may provide insight into potential novel therapeutic avenues that could be acutely initiated depending on time of injury.

## Acknowledgements

This work was supported by Kentucky Spinal Cord and Head Injury Research Trust (contract# 18-2), NS108529-01, NS114404, Norton Healthcare, and the Commonwealth of Kentucky Challenge for Excellence. We thank Christine Yarberry for surgical assistance, Johnny Morehouse and Jason Beare for behavioral analyses.

## Supplementary Information

**Supplementary Methods.** *<u>Spinal cord injury and post-surgery animal care</u>.*

**Supplementary Table S1.** Contusion parameters for each individual animal

**Supplementary Table S2.** qPCR primers

**Supplementary Table S3**. Results of RM ANOVA for BMS and ladder test

**Supplementary Figures S1 and S2. Circadian oscillations of selected clock pathway mRNAs in the brain stem, the cerebellum and the liver of C57Bl6 mice.** The data are from the publicly available circadian transcriptome database (http://circadb.hogeneschlab.org/mouse). All presented mRNAs show significant circadian oscillations in all three tissues (JTK p<0.05) except *Cry1* (non-significant cycling in the brain stem /JTK p=0.058/) and *B2m* (no cycling in any tissue /JTK p=1/).

## References

1. Durgan DJ, Young ME. The cardiomyocyte circadian clock: emerging roles in health and disease. Circ Res. 2010;106(4):647–58. Epub 2010/03/06. doi: 10.1161/CIRCRESAHA.109.209957. PubMed PMID: 20203314; PubMed Central PMCID: PMCPMC3223121.

2. Traverse JH. Of mice and men: the quest to determine a circadian basis for myocardial protection in ischemia/reperfusion injury. Circ Res. 2013;112(10):e115–7. Epub 2013/05/11. doi: 10.1161/CIRCRESAHA.113.301079. PubMed PMID: 23661714.

3. Shokri HM, El Nahas NM, Aref HM, Dawood NL, Abushady EM, Abd Eldayem EH, et al. Factors related to time of stroke onset versus time of hospital arrival: A SITS registry-based study in an Egyptian stroke center. PLoS One. 2020;15(9):e0238305. Epub 2020/09/12. doi: 10.1371/journal.pone.0238305. PubMed PMID: 32915811; PubMed Central PMCID: PMCPMC7485782.

4. Musiek ES, Holtzman DM. Mechanisms linking circadian clocks, sleep, and neurodegeneration. Science. 2016;354(6315):1004–8. Epub 2016/11/26. doi: tzman PubMed PMID: 27885006; PubMed Central PMCID: PMC5219881.

5. Carroll RG, Timmons GA, Cervantes-Silva MP, Kennedy OD, Curtis AM. Immunometabolism around the Clock. Trends in molecular medicine. 2019;25(7):612–25. Epub 2019/06/04. doi: 10.1016/j.molmed.2019.04.013. PubMed PMID: 31153819.

6. Cuddapah VA, Zhang SL, Sehgal A. Regulation of the Blood-Brain Barrier by Circadian Rhythms and Sleep. Trends Neurosci. 2019;42(7):500–10. Epub 2019/06/30. doi: 10.1016/j.tins.2019.05.001. PubMed PMID: 31253251; PubMed Central PMCID: PMCPMC6602072.

7. Lowrey PL, Takahashi JS. Genetics of circadian rhythms in Mammalian model organisms. Advances in genetics. 2011;74:175–230. Epub 2011/09/20. doi: 10.1016/B978-0-12-387690-4.00006-4. PubMed PMID: 21924978; PubMed Central PMCID: PMC3709251.

8. Zhang R, Lahens NF, Ballance HI, Hughes ME, Hogenesch JB. A circadian gene expression atlas in mammals: implications for biology and medicine. Proceedings of the National Academy of Sciences of the United States of America. 2014;111(45):16219–24. Epub 2014/10/29. doi: 10.1073/pnas.1408886111. PubMed PMID: 25349387; PubMed Central PMCID: PMC4234565.

9. Hor CN, Yeung J, Jan M, Emmenegger Y, Hubbard J, Xenarios I, et al. Sleepwake-driven and circadian contributions to daily rhythms in gene expression and chromatin accessibility in the murine cortex. Proceedings of the National Academy of Sciences of the United States of America. 2019;116(51):25773–83. Epub 2019/11/30. doi: 10.1073/pnas.1910590116. PubMed PMID: 31776259; PubMed Central PMCID: PMCPMC6925978.

10. Yang N, Smyllie NJ, Morris H, Goncalves CF, Dudek M, Pathiranage DRJ, et al. Quantitative live imaging of Venus::BMAL1 in a mouse model reveals complex dynamics of the master circadian clock regulator. PLoS Genet. 2020;16(4):e1008729. Epub 2020/05/01. doi: 10.1371/journal.pgen.1008729. PubMed PMID: 32352975; PubMed Central PMCID: PMCPMC7217492.

11. Durgan DJ, Pulinilkunnil T, Villegas-Montoya C, Garvey ME, Frangogiannis NG, Michael LH, et al. Short communication: ischemia/reperfusion tolerance is time-of-day-dependent: mediation by the cardiomyocyte circadian clock. Circ Res. 2010;106(3):546–50. Epub 2009/12/17. doi: 10.1161/CIRCRESAHA.109.209346. PubMed PMID: 20007913; PubMed Central PMCID: PMCPMC3021132.

12. Nguyen KD, Fentress SJ, Qiu Y, Yun K, Cox JS, Chawla A. Circadian gene Bmal1 regulates diurnal oscillations of Ly6C(hi) inflammatory monocytes. Science. 2013;341(6153):1483–8. Epub 2013/08/24. doi: 10.1126/science.1240636. PubMed PMID: 23970558; PubMed Central PMCID: PMCPMC3836670.

13. Deng W, Zhu S, Zeng L, Liu J, Kang R, Yang M, et al. The Circadian Clock Controls Immune Checkpoint Pathway in Sepsis. Cell reports. 2018;24(2):366–78. Epub 2018/07/12. doi: 10.1016/j.celrep.2018.06.026. PubMed PMID: 29996098; PubMed Central PMCID: PMCPMC6094382.

14. Curtis AM, Fagundes CT, Yang G, Palsson-McDermott EM, Wochal P, McGettrick AF, et al. Circadian control of innate immunity in macrophages by miR-155 targeting Bmal1. Proceedings of the National Academy of Sciences of the United States of America. 2015;112(23):7231–6. Epub 2015/05/23. doi: 10.1073/pnas.1501327112. PubMed PMID: 25995365; PubMed Central PMCID: PMCPMC4466714.

15. Sutton CE, Finlay CM, Raverdeau M, Early JO, DeCourcey J, Zaslona Z, et al. Loss of the molecular clock in myeloid cells exacerbates T cell-mediated CNS autoimmune disease. Nature communications. 2017;8(1):1923. Epub 2017/12/14. doi: 10.1038/s41467-017-02111-0. PubMed PMID: 29234010; PubMed Central PMCID: PMCPMC5727202.

16. Vinall PE, Kramer MS, Heinel LA, Rosenwasser RH. Temporal changes in sensitivity of rats to cerebral ischemic insult. J Neurosurg. 2000;93(1):82–9. Epub 2000/07/07. doi: 10.3171/jns.2000.93.1.0082. PubMed PMID: 10883909.

17. Tischkau SA, Cohen JA, Stark JT, Gross DR, Bottum KM. Time-of-day affects expression of hippocampal markers for ischemic damage induced by global ischemia. Experimental neurology. 2007;208(2):314–22. Epub 2007/10/16. doi: 10.1016/j.expneurol.2007.09.003. PubMed PMID: 17936274.

18. Martinez-Vargas M, Gonzalez-Rivera R, Soto-Nunez M, Cisneros-Martinez M, Huerta-Saquero A, Morales-Gomez J, et al. Recovery after a traumatic brain injury depends on diurnal variations effect of cystatin C. Neurosci Lett. 2006;400(1-2):21–4. Epub 2006/03/08. doi: 10.1016/j.neulet.2006.02.010. PubMed PMID: 16519999.

19. Weil ZM, Karelina K, Su AJ, Barker JM, Norman GJ, Zhang N, et al. Time-of-day determines neuronal damage and mortality after cardiac arrest. Neurobiol Dis. 2009;36(2):352–60. Epub 2009/08/12. doi: 10.1016/j.nbd.2009.07.032. PubMed PMID: 19664712; PubMed Central PMCID: PMCPMC2760634.

20. Schallner N, Lieberum JL, Gallo D, LeBlanc RH, 3rd, Fuller PM, Hanafy KA, et al. Carbon Monoxide Preserves Circadian Rhythm to Reduce the Severity of Subarachnoid Hemorrhage in Mice. Stroke. 2017;48(9):2565–73. Epub 2017/07/28. doi: 10.1161/STROKEAHA.116.016165. PubMed PMID: 28747460; PubMed Central PMCID: PMCPMC5575974.

21. Beker MC, Caglayan B, Yalcin E, Caglayan AB, Turkseven S, Gurel B, et al. Time-of-Day Dependent Neuronal Injury After Ischemic Stroke: Implication of Circadian Clock Transcriptional Factor Bmal1 and Survival Kinase AKT. Mol Neurobiol. 2018;55(3):2565–76. Epub 2017/04/20. doi: 10.1007/s12035-017-0524-4. PubMed PMID: 28421530.

22. Slomnicki LP, Myers SA, Saraswat Ohri S, Parsh MV, Andres KR, Chariker JH, et al. Improved locomotor recovery after contusive spinal cord injury in Bmal1(-/-) mice is associated with protection of the blood spinal cord barrier. Scientific reports. 2020;10(1):14212. Epub 2020/08/28. doi: 10.1038/s41598-020-71131-6. PubMed PMID: 32848194; PubMed Central PMCID: PMCPMC7450087.

23. Ohri SS, Maddie MA, Zhao Y, Qiu MS, Hetman M, Whittemore SR. Attenuating the endoplasmic reticulum stress response improves functional recovery after spinal cord injury. Glia. 2011;59(10):1489–502. Epub 2011/06/04. doi: 10.1002/glia.21191. PubMed PMID: 21638341; PubMed Central PMCID: PMC3391751.

24. Basso DM, Fisher LC, Anderson AJ, Jakeman LB, McTigue DM, Popovich PG. Basso Mouse Scale for locomotion detects differences in recovery after spinal cord injury in five common mouse strains. J Neurotrauma. 2006;23(5):635–59. Epub 2006/05/13. doi: 10.1089/neu.2006.23.635. PubMed PMID: 16689667.

25. Kim JH, Song SK, Burke DA, Magnuson DS. Comprehensive locomotor outcomes correlate to hyperacute diffusion tensor measures after spinal cord injury in the adult rat. Experimental neurology. 2012;235(1):188–96. Epub 2011/11/29. doi: 10.1016/j.expneurol.2011.11.015. PubMed PMID: 22119625; PubMed Central PMCID: PMC3334428.

26. Gaudet AD, Fonken LK, Ayala MT, Bateman EM, Schleicher WE, Smith EJ, et al. Spinal Cord Injury in Rats Disrupts the Circadian System. eNeuro. 2018;5(6). Epub 2019/01/11. doi: 10.1523/ENEURO.0328-18.2018. PubMed PMID: 30627655; PubMed Central PMCID: PMCPMC6325559.

27. Magnuson DS, Trinder TC, Zhang YP, Burke D, Morassutti DJ, Shields CB. Comparing deficits following excitotoxic and contusion injuries in the thoracic and lumbar spinal cord of the adult rat. Experimental neurology. 1999;156(1):191–204. Epub 1999/04/08. doi: 10.1006/exnr.1999.7016. PubMed PMID: 10192790.

28. Hughes ME, Abruzzi KC, Allada R, Anafi R, Arpat AB, Asher G, et al. Guidelines for Genome-Scale Analysis of Biological Rhythms. J Biol Rhythms. 2017;32(5):380–93. Epub 2017/11/04. doi: 10.1177/0748730417728663. PubMed PMID: 29098954; PubMed Central PMCID: PMCPMC5692188.

29. Ohri SS, Hetman M, Whittemore SR. Restoring endoplasmic reticulum homeostasis improves functional recovery after spinal cord injury. Neurobiol Dis. 2013;58:29–37. Epub 2013/05/11. doi: 10.1016/j.nbd.2013.04.021. PubMed PMID: 23659896; PubMed Central PMCID: PMC3748169.

30. Myers SA, Andres KR, Hagg T, Whittemore SR. CD36 deletion improves recovery from spinal cord injury. Experimental neurology. 2014;256:25–38. Epub 2014/04/03. doi: 10.1016/j.expneurol.2014.03.016. PubMed PMID: 24690303; PubMed Central PMCID: PMC4086463.

31. Saraswat Ohri S, Bankston AN, Mullins SA, Liu Y, Andres KR, Beare JE, et al. Blocking Autophagy in Oligodendrocytes Limits Functional Recovery after Spinal Cord Injury. J Neurosci. 2018;38(26):5900–12. Epub 2018/05/26. doi: 10.1523/JNEUROSCI.0679-17.2018. PubMed PMID: 29793971; PubMed Central PMCID: PMC6021994.

32. Saraswat Ohri S, Burke DA, Andres KR, Hetman M, Whittemore SR. Acute Neural and Proteostasis Messenger Ribonucleic Acid Levels Predict Chronic Locomotor Recovery after Contusive Spinal Cord Injury. J Neurotrauma. 2020. Epub 2020/10/21. doi: 10.1089/neu.2020.7258. PubMed PMID: 33076743.

33. Musiek ES, Lim MM, Yang G, Bauer AQ, Qi L, Lee Y, et al. Circadian clock proteins regulate neuronal redox homeostasis and neurodegeneration. J Clin Invest. 2013;123(12):5389–400. Epub 2013/11/26. doi: 10.1172/JCI70317. PubMed PMID: 24270424; PubMed Central PMCID: PMC3859381.

34. Yang G, Chen L, Grant GR, Paschos G, Song WL, Musiek ES, et al. Timing of expression of the core clock gene Bmal1 influences its effects on aging and survival. Science translational medicine. 2016;8(324):324ra16. Epub 2016/02/05. doi: 10.1126/scitranslmed.aad3305. PubMed PMID: 26843191; PubMed Central PMCID: PMC4870001.

35. Imai R, Makino H, Katoh T, Kimura T, Kurita T, Hokamura K, et al. Desflurane anesthesia shifts the circadian rhythm phase depending on the time of day of anesthesia. Scientific reports. 2020;10(1):18273. Epub 2020/10/28. doi: 10.1038/s41598-020-75434-6. PubMed PMID: 33106509; PubMed Central PMCID: PMCPMC7588451.

36. Jirkof P, Tourvieille A, Cinelli P, Arras M. Buprenorphine for pain relief in mice: repeated injections vs sustained-release depot formulation. Lab Anim. 2015;49(3):177–87. Epub 2014/12/10. doi: 10.1177/0023677214562849. PubMed PMID: 25488320.

37. Geisler FH, Coleman WP, Grieco G, Poonian D, Sygen Study G. Measurements and recovery patterns in a multicenter study of acute spinal cord injury. Spine. 2001;26(24 Suppl):S68–86. Epub 2002/01/24. doi: 10.1097/00007632-200112151-00014. PubMed PMID: 11805613.

38. Steeves JD, Lammertse D, Curt A, Fawcett JW, Tuszynski MH, Ditunno JF, et al. Guidelines for the conduct of clinical trials for spinal cord injury (SCI) as developed by the ICCP panel: clinical trial outcome measures. Spinal cord. 2007;45(3):206–21. Epub 2006/12/21. doi: 10.1038/sj.sc.3102008. PubMed PMID: 17179972.

39. Zhang Y, Guan Z, Reader B, Shawler T, Mandrekar-Colucci S, Huang K, et al. Autonomic dysreflexia causes chronic immune suppression after spinal cord injury. J Neurosci. 2013;33(32):12970–81. Epub 2013/08/09. doi: 10.1523/JNEUROSCI.1974-13.2013. PubMed PMID: 23926252; PubMed Central PMCID: PMC3735880.

